# AI agent supervision of structural model building and refinement

**DOI:** 10.64898/2026.10.05.756902

**Authors:** Kuen-Phon Wu, Chuan-Kai Chueh

## Abstract

Structural model building and refinement need many decisions between different programs. These decisions often depend on an experienced researcher who knows how the programs work together and how to read experimental data. We used Claude Code with Claude Opus 5.5 to supervise this work for 24 cryo-EM and X-ray cases. The work finished in about 36 hours on one workstation. The agent prepared the inputs, ran established programs, read the validation results, compared candidate models and chose the next step. For 15 deposited cryo-EM models with weak starting geometry, the median MolProbity score improved from 2.62 to 1.76. For three crystal structures, the agent models came close to the published R_free_ values. Chemical information such as ligands, ions and modifications must be provided by the researcher, in the same way as the sequence. With this information, the agent can place and check these components. These results show that agent supervision is acceptable when every decision can be inspected. The researcher makes the final scientific judgment. We suggest practical requirements for this kind of use. Future AI agents may help researchers obtain high quality and reliable structures of biological macromolecules.

## The work between programs

Protein structure prediction and automated model building have changed the starting point of structural biology. AlphaFold 3 predicts structures of molecular complexes ^1^. ModelAngelo builds protein models from cryo-EM maps and sequence information ^2^. These tools do not remove the need to examine an experimental model after it has been built. The model must fit the measured data, maintain plausible chemistry and represent uncertainty honestly. Side chains, interfaces, waters, ligands and ions may each need separate attention. Structural biologists already have capable programs for these tasks. Phenix supports refinement and validation ^3^. Coot supports visualization and local building ^4^. MolProbity, CaBLAM and map-based measures reveal different problems ^5-7^. Phenix ^8^, CCP4 ^9^, Phaser ^10^, Buccaner ^11^, and/or ARP/wARP ^12^ are needed help the X-ray structure determination. For many projects, the bottleneck is not the absence or a choice of a program. It is the work between programs. For examples, a user decides what to run, interprets a report, sees a new problem, changes the settings and decides whether to keep the result. A model may fit a map with an improved model-to-map correlations while acquiring more clashes. A better rotameric distributions might pull the backbone away from density. A water search can add plausible and implausible sites together.

Making decisions have often depended on an experienced researcher who knows how the programs interact and how to interpret experimental maps. We propose that an advanced AI agent can now perform much of this supervisory work in an efficient timing. The agent does not calculate a density map or optimize atomic coordinates by itself. It uses established programs, checks their output and selects the next step. In this role, an agent is closer to a workflow supervisor than to a new refinement engine to avoid AI-lead hallucination. The structural biologist still decides whether the final model is scientifically acceptable.

Our experience provides a concrete demonstration of this idea. Anthropic’s Claude Code, using advanced Claude Opus 5.5 (released on September 22, 2026) at the high effort setting, coordinated model building, refinement and validation across 24 cryo-EM and X-ray cases. The computational work finished over about 36 hours on one workstation. We did not test earlier or smaller language models. We therefore do not claim that every AI agent can do this. The result shows what one newly released agent was able to do under investigator guidance at the time of this study. The wider goal is that this supervisory role may become a normal part of structural biology as capable agents become more available.

### What agent supervision means

Supervision begins when the investigator supplies an experimental task and the relevant data. For cryo-EM, these may include a full map, half-maps, sequence information and an initial model. For X-ray crystallography, they may include diffraction data, free-reflection flags, sequence and a search or starting model. The investigator also defines the aim. The aim may be to build a model from a map, find defects in a deposited model or improve a structure already refined by the laboratory. When nonprotein components are relevant, the investigator should also provide the known sample composition, expected ligands or cofactors, likely ions and post-translational modification information. These are experimental facts that cannot be recovered reliably from coordinates or density alone. **Figure 1** shows the division of work.

**Figure 1.**
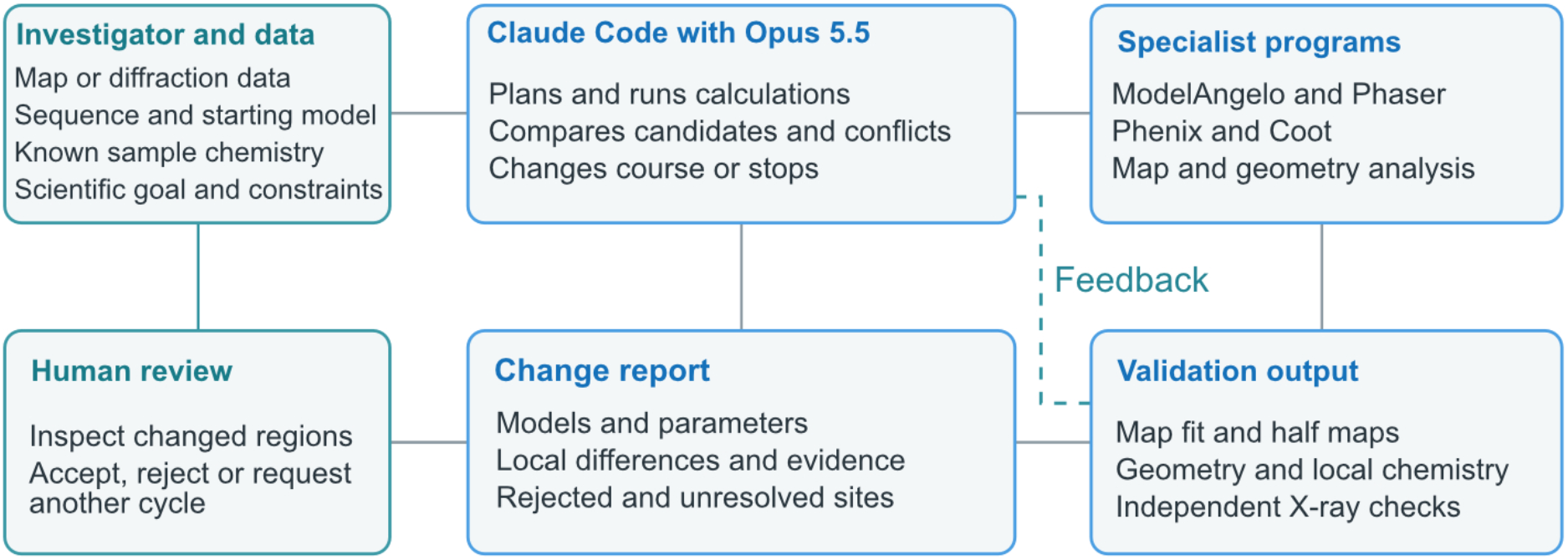
AI agent supervision across structural biology programs. The investigator supplies experimental data, known sample chemistry and a goal. Claude Code with Claude Opus 5.5 prepares and runs specialist programs. It reads validation output, compares candidate models and chooses another calculation or an abstention when criteria are not met. The investigator reviews the proposed structural changes and can accept, reject or request another cycle. The diagram describes the tested division of work, not a claim that every AI agent can perform it. Nonprotein chemical assignment was not tested in the de novo cases.

The agent prepares inputs, runs the specialist programs and reads their outputs. It can compare several candidate models rather than accept the latest file. It can change a weight, constrain a local operation, return to an earlier model or ask for another calculation. It can write scripts to connect programs that do not share a single interface. After each cycle, it should record the program version, command, parameters, candidate model, validation values and reason for the next step. This record matters because the same final score could be reached by different routes.

The word supervision does not mean that the agent is an independent authority on structural chemistry. It means that the agent manages a sequence of calculations and checks whether each result meets stated criteria. An agent can detect that a candidate has a worse clashscore or weaker map agreement. It can recognize that two metrics disagree and direct attention to the affected region. It cannot make a weak density feature unambiguous by describing it confidently. Its recommendations must stay linked to the experimental evidence and remain open to rejection by the investigator.

This role differs from a fixed pipeline. A fixed pipeline follows a predetermined sequence and may apply the same settings even when an intermediate result is poor. Agent supervision allows the route to change in response to the result. It is also different from asking a chatbot for advice while manually operating each program. The agent performs the calculations, compares the resulting files and prepares a focused report for review. The practical value lies in the connection of planning, execution, validation and revision.

### An experience across 24 cases

Our laboratory explored this approach through several working sessions with Claude Code and Claude Opus 5.5. The investigator gave the initial goals, chose or requested additional cases, and later asked for closer checks of particular outputs. The agent obtained files, prepared scripts, ran structural programs, compared models and reported outcomes. This was not an unattended one-prompt experiment. It was an interactive project in which the agent carried out most of the repeated computational work while the investigator continued to guide the scientific questions.

The exercise included 15 deposited cryo-EM models for re-refinement. Five further cryo-EM cases began with maps and sequences. ModelAngelo supplied the initial protein models in those five cases, after which the agent prepared, corrected, refined and validated the output. Four in-house cases extended the exercise where one used cryo-EM data and three used high-resolution X-ray data. The cohorts, entries, software and selection rules are documented in **Supplementary Tables S1–S4**. The work was completed in about 36 hours of elapsed project time on one Ubuntu workstation with a 16-core AMD CPU, 128 GB RAM and an NVIDIA RTX 4070 Ti SUPER GPU. Per-case runtime, investigator time and token use were not measured separately. The 36-hour observation is therefore a measure of project throughput, not a controlled speed comparison with human researchers.

In the 15 cryo-EM re-refinement cases, median MolProbity score improved from 2.62 to 1.76 and median clashscore fell from about 21 to 6. Median all-atom Q-score increased from 0.531 to 0.568 ^6^. Selected models kept masked map correlation within 0.01 of the deposited value or improved it. The Cα root-mean-square displacement from the deposited model ranged from 0.10 to 0.66 Å. **Figure 2** summarize the geometry and Q-score changes as well as examples of structures of depositions and AI-supervision. Per-entry values are in **Supplementary Tables S5–S8**. These results are consistent with many local corrections rather than broad changes in protein fold. In low-resolution cases, the agent removed some weakly supported residues or side-chain atoms. Those changes reduce unsupported modelling but do not prove the removed atoms have no physical position.

**Figure 2.**
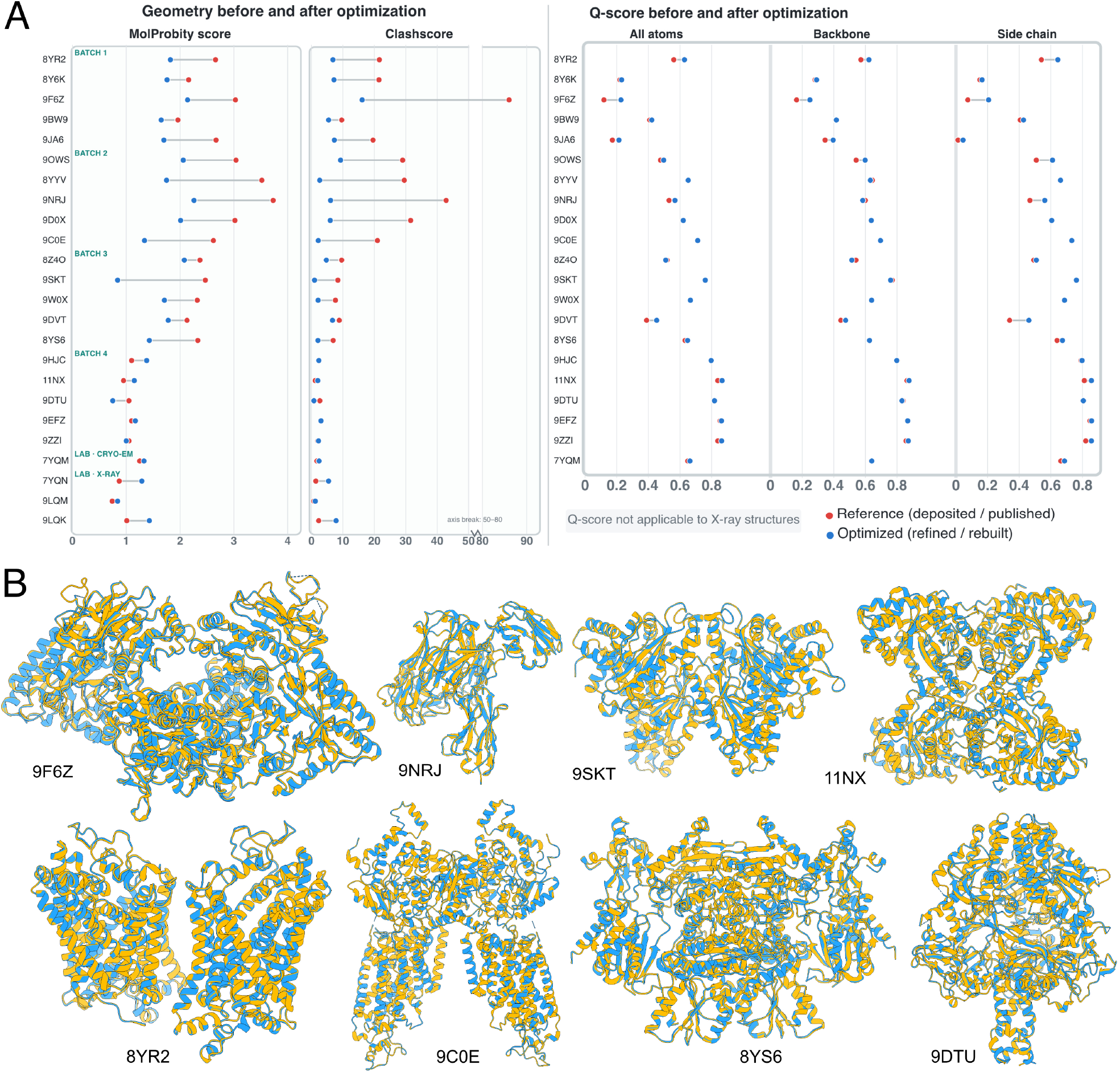
Evidence from cryo-EM and X-ray workflows. (A)Metrics of geometry of paired MolProbity and clashscores for 21 deposited cryo-EM models before and after agent-supervised re-refinement. Each line represents one case. Median all-atom, backbone and side-chain Q-scores measured against the same deposited primary maps. Red indicates deposited models and blue indicates selected agent-guided models. Per-entry values for panels A and B are in Supplementary Tables. (B) Representative overlays of eight selected cases show that the AI-guided refined models (yellow) remain consistent with the original models (dodger blue), with no obvious refinement-induced artifacts or major structural distortions.

The five map-and-sequence cases demonstrate a different type of workflow. ModelAngelo built protein coordinates from the map and sequence. The agent then refined the candidate structures and checked their quality. The final protein models covered 95.5 to 100% of residues in the corresponding published coordinate models. Matched Cα positions differed by 0.10 to 0.21 Å. The Q-scores and model comparisons are in **Supplementary Tables S9 and S11**. This supports the feasibility of connecting automated building to supervised refinement. It does not mean the language model inferred atomic coordinates directly from density. This exercise tested protein building, not complete chemical assignment. Ligands, ions and glycans were outside its scope and were not built. The deposited 9DTU model contains glycans ^13^. The deposited 9HJC (https://www.rcsb.org/structure/9HJC) model contains thiamine (VIB) and magnesium. The deposited 9EFZ ^14^ and 9ZZI ^15^ models also contain nonprotein components. Their absence from our de novo candidates is a limitation of the input and workflow, not evidence that those components are absent from the samples.

The in-house cases added a useful check on the meaning of success. For cryo-EM structure 7YQM ^16^, some candidate models improved map-based measures but failed the geometry acceptance rule. The deposited model was retained (**Supplementary Tables S12 and S13**). In the three X-ray cases, the agent-guided models were built from diffraction data without using the published coordinates during the work. A second refinement pass, requested by the investigator, improved local fit and water placement. Final hydrogen-free R_free_ values were 0.182, 0.207 and 0.238. The published models gave 0.173, 0.197 and 0.229 on the same data. Refinement rules, stepwise results and model composition are in **Supplementary Tables S14–S18**.

The agent-guided models approached, but did not exceed, the published values. They show that a long multi-program route was feasible. They do not show that the agent replaced expert final refinement.

### The important result is a change of course

The global improvements are useful, but they are not the central observation of this study. The most informative moments came when the agent changed the workflow after inspecting an intermediate result. During a map-weight scan, a higher map correlation came with more clashes. The agent moved toward automatic weighting and applied geometry checks before choosing a candidate. In another part of the work, Coot side-chain fitting shifted backbone atoms. The workflow was revised to transfer supported side-chain torsions while retaining the original backbone. When B-factor refinement or atomic displacement parameter (ADP) refinement ^17^ weakened map agreement, the agent compared a branch that kept the deposited B factors. **Table 1** lists these examples and the associated human checks. Further case-specific problems and responses are documented in **Supplementary Table S19**.

**Table 1.** Examples of adaptive decisions during the working sessions. The table links an observed problem to the agent’s change of course and to the local evidence that still requires investigator review. It is an interpretive summary of the recorded workflow, not a claim that every proposed correction was accepted.

| Observed problem | Agent response | Investigator review |
| --- | --- | --- |
| Higher map correlation also raises clashes | Test automatic weighting and apply geometry checks before model selection. | Inspect affected atoms against local density and peptide geometry. |
| Side-chain fitting shifts the backbone | Retain supported side-chain torsions but restore the starting backbone. | Check local side-chain density and backbone continuity. |
| B-factor refinement weakens map agreement | Compare a second branch that preserves deposited B factors. | Review local density and the same map-fit measure for both branches. |
| Water search produces too many sites | Filter by density, nearby N or O contacts and short-contact exclusion. | Check whether each site is water, ion, ligand or unsupported. |
| No candidate meets acceptance rules | Keep the starting model and report the failed criteria. | Decide whether manual rebuilding or further data analysis is needed. |

These responses resemble decisions made by an experienced structural biologist. They require more than knowing one command. The user must understand that the outputs of different programs can conflict and that a locally improved number can harm another part of the model. Our cases show that one AI agent could notice several such conflicts and respond operationally. They do not establish that the agent would recognize every error or choose the best response in every new structure.

Water placement illustrates the same distinction. Automated searching can return more candidates than the data support. In the map-and-sequence work, proposed waters were filtered by map support, a plausible contact to a nitrogen or oxygen atom within 3.5 Å, and absence of a very short contact below 2.2 Å. Even a candidate that passes these filters may actually be an ion, a ligand atom or a feature of an unbuilt component. An agent can narrow the list and flag uncertainty. The assignment of chemical identity still requires local interpretation. The same care applies to metal sites ^18^.

The de novo cases make this limitation concrete. Protein coverage, Cα agreement, Q-score and geometry cannot establish that a model has the correct glycans, ligands or ions. In a post hoc comparison, three, eight and 17 candidate waters in 9HJC, 9DTU ^13^ and 9EFZ ^14^ occupied sites assigned to ligands or ions in the published models (**Supplementary Table S11**). A strong density feature may suggest a site, but density alone may not identify the chemical species. The agent could ask the investigator for a candidate list and restraint information, test each plausible model against local density and chemical environment, and report sites that remain unexplained. The investigator should confirm assignments using the experimental composition and biological context. If the evidence does not distinguish candidates, the site should remain unresolved rather than receive a confident but unsupported label.

A useful agent output is therefore not only a refined coordinate file. It is a change report. For each altered region, the report should identify the original and proposed coordinates, the operation used, the local fit to the data, the geometry change and any new concern. It should name candidates that were rejected and regions that remain unresolved. The decision to keep the original 7YQM model is as important to this argument as a case with a lower clashscore. Supervision includes knowing when the available operations did not produce a defensible improvement.

### Trust is an operational property

For an agent-supervised workflow, trust should mean that its decisions can be inspected and challenged. A good global score is not enough. Clashscore, Ramachandran and rotamer analysis, peptide geometry and CaBLAM ^19^ examine different aspects of coordinate quality. Map correlation, Q-score and local density inspect agreement with cryo-EM data. Half-map tests can reveal some forms of overfitting. The case-level half-map results are in **Supplementary Table S10**. For X-ray structures, R_free_ and local real-space measures add independent checks. Community validation work has shown that geometry and fit to experimental data are not interchangeable ^5,20^.

Our own results show why these checks must disagree sometimes. CaBLAM outlier rates increased in seven of the 15 cryo-EM re-refinement cases, even as other global measures improved. One ligand had weaker local map agreement after refinement. Neither result can be dismissed by the improved median MolProbity score ^7^. The agent’s supervisory role is valuable when it makes these conflicts visible and returns them to the scientist for review. The agent should also preserve a route back to the original or previous model. A new candidate should be written to a separate work area. A change should not be accepted solely because it is newer, visually attractive or supported by one score. The investigator should inspect every altered backbone segment and every change to an interface, ligand, ion or water. If local evidence is weak, keeping the original model or leaving the site unresolved is a valid outcome. Such review is not a ceremonial last step. It is part of the method.

This division of responsibility need not make the workflow slow. The agent can perform the broad search, run validation repeatedly and present a short list of consequential changes. The investigator then spends time on the regions where judgment matters. This may be especially useful in laboratories that solve structures only occasionally. The agent can help them access the full capability of existing software and see why a particular correction was proposed.

### What future agents may change

This study tested AI agent configuration. It did not compare Claude Opus 5.5 with earlier versions, other vendors or a fixed automated pipeline. The present results should not be presented as a property of all language models. They provide a dated example of a capability that can now be examined directly. Later agents may handle longer projects, diagnose more types of error and require fewer interventions. These are reasonable directions to test, not findings established by the present cases.

The most useful next comparison would hold the structural tasks and acceptance rules constant while changing the agent. It should record elapsed time, human intervention time, tool calls, token use, failed branches and computational cost. It should also ask independent structural biologists to review the changed regions without knowing which model produced them. Local correctness, evidence for water and ion assignments, and the ability to abstain should matter more than a single global score. This design would show whether a newer agent is better at supervision rather than merely more fluent at reporting its work. It should include cases in which the investigator supplies chemical information and cases in which that information is incomplete. This would test when the agent can make a justified assignment and when it must ask for help.

More complete automation will require progress on both sides of the interface. Structural software needs better ways to propose and test chemically specified ligands, glycans and ions, to compare them with water or an unassigned feature, and to expose local evidence in a form an agent can use. Future agents need to combine that evidence with sample preparation, sequence annotation and investigator-supplied knowledge. They also need to recognize when key information is missing. An agent may eventually suggest a chemical identity with strong evidence, but it should not silently turn a plausible guess into a deposited atom assignment. The present benchmark does not test that future capability.

The approach is not tied to a permanent software stack. ModelAngelo (https://github.com/3dem/model-angelo) and Coot (https://www2.mrc-lmb.cam.ac.uk/personal/pemsley/coot/) are publicly available, while Phenix (https://phenix-online.org/license) is available without cost for non-profit research under its own license. Laboratories can choose different tools as long as inputs, versions, commands and outputs are recorded. A general agent can connect the tools already used by a laboratory. Dedicated systems may provide further advantages, and related agent-based efforts are emerging ^21^. The scientific principle is broader than one product. Let the agents supervise the transitions between established programs, then judge their work against the experimental data.

### A route for routine use

A laboratory can begin with a modest task. The researcher first builds or refines a model using their normal workflow. They then provide the agent with the data, current model, known sample chemistry and a request to find remaining defects. If a ligand, ion or glycan is expected, the researcher should identify it or provide plausible candidates and any available chemical restraints. The agent should keep the original files unchanged, make candidates in a separate directory, run more than one type of validation and explain why it selected or rejected each candidate. The final output should include coordinates, metrics, commands, a residue-level change list and a list of unassigned features. The researcher inspects the listed sites and either accepts, rejects or requests another cycle.

The same supervisory pattern can be used earlier in structure determination. For cryo-EM, the agent can connect map preparation, automated model building, refinement and validation. For X-ray crystallography, it can connect molecular replacement, refinement, solvent modelling and local rebuilding. The programs and data differ, but the supervisory questions remain familiar. Did the last operation improve the intended region? Did it create a new problem elsewhere? Is the proposed chemistry plausible? Which regions still lack convincing evidence?

Our 24-case experience suggests that the AI agent (Claude Opus 5.5) can already carry out many of these iterative decisions at a useful pace. It does not remove the need for structural biology expertise. It changes where that expertise is applied. Researchers can direct the scientific goal and examine consequential local changes, while the agent supervises calculations, detects conflicts and prepares the evidence for review. As AI agents become more widely used in structural biology, we encourage explicit disclosure of their role and documentation of the main computational decisions. Relevant prompts, execution records and model comparisons can help investigators assess proposed changes and allow others to understand how the final model was selected. Future agents may extend this role. Their progress should be measured by how well they support correct, inspectable structural decisions, not by how confidently they claim to have finished a model.

### Cases with PDB entry IDs and available citations

Cryo-EM re-refinement. 8YR2 ^22^, 8Y6K ^23^, 9F6Z ^24^, 9BW9 ^25^, 9JA6 ^26^, 9OWS ^27^, 8YYV ^28^, 9NRJ ^29^, 9D0X ^30^, 9C0E ^31^, 8Z4O, 9SKT ^32^, 9W0X, 9DVT ^33^ and 8YS6 ^34^.

Cryo-EM map-and-sequence cases. 9HJC, 11NX ^35^, 9DTU ^13^, 9EFZ ^14^ and 9ZZI ^15^.

In-house cases. 7YQM ^16^, 7YQN ^16^, 9LQM ^36^ and 9LQK ^36^.

## Acknowledgements

K.P.W. is supported by Academia Sinica (AS) intramural funding and grants (AS-IV-115-L01 and AS-IAIA-114-L02) and by a grant from Taiwan’s National Science and Technology Council (NSTC-115-2113-M-001-004-). C.K.C. acknowledges support from the NSTC Graduate Research Fellowship.

## Declaration of generative AI and AI-assisted technologies in the manuscript preparation process

During the preparation of this work, the author used ChatGPT and Claude to revise English grammar and identify typographical errors, mislabels, and other editing errors. The author reviewed and edited the outputs as needed and take full responsibility for the content of the published article.

