## Supplementary tables S1-S19 for "AI agent supervision of structural model building and refinement"

**Table of contents:**

**Supplementary Table S1. Exercise selection**

**Supplementary Table S2. Cryo-EM entries and deposited validation**

**Supplementary Table S3. Software used in the exercise**

**Supplementary Table S4. Cryo-EM candidate selection rules**

**Supplementary Table S5. Batch 1 geometry and map-fit metrics**

**Supplementary Table S6. Batch 2 geometry and map-fit metrics**

**Supplementary Table S7. Batch 3 geometry and map-fit metrics**

**Supplementary Table S8. Q-scores for re-refined cryo-EM entries**

**Supplementary Table S9. Q-scores for de novo models and published models**

**Supplementary Table S10. Half-map cross-validation results**

**Supplementary Table S11. De novo versus published model metrics**

**Supplementary Table S12. In-house entry metadata**

**Supplementary Table S13. Cryo-EM re-refinement of 7YQM**

**Supplementary Table S14. Crystal structure decision rules**

**Supplementary Table S15. Crystal first-pass final and published metrics**

**Supplementary Table S16. Crystal second-pass refinement steps**

**Supplementary Table S17. Crystal parameter tests and decisions**

**Supplementary Table S18. Crystal model composition compared with published entries**

**Supplementary Table S19. Special handling and identified problems**

### Study design and methods

#### Supplementary Table S1. Exercise selection

The original report describes four batches. Its batch 4 comprises the five de novo cryo-EM cases.

| Batch | Goal | Selection criteria | Pool | Picked |
|---|---|---|---|---|
| 1 | Re-refine user-chosen poor models | User list: 8YR2, 9D0K, 8Y6K, 9F6Z, 9BW9. 9D0K was withdrawn from the PDB (status WDRN) and replaced by 9JA6 at the user's request. | — | 8YR2, 8Y6K, 9F6Z, 9BW9, 9JA6 |
| 2 | Poor recent models with nucleic acid, protein–protein or drug | Released ≥ 2025-01-01 · 2–4 Å · ≤ 300 kDa · clashscore ≥ 20. Ranked by clashscore >20 + Rama outliers >0.5 % + rotamer outliers > 3 %. A drug-like ligand was required for the ligand category. | 29 NA<br>79 PPI<br>87 ligand | 9OWS, 8YYV, 9NRJ, 9D0X, 9C0E |
| 3 | Moderately good recent models | Released ≥ 2025 · 150–350 kDa · 4 < clashscore < 10 · rotamer outliers ≥ 2 % · Rama outliers ≥ 0.3 % · 2–4 Å. Chosen for diversity: glycoprotein, metal site, membrane protein, protein–DNA, drug + Fe–S clusters. | 47 | 8Z4O, 9SKT, 9W0X, 9DVT, 8YS6 |
| 4 | Rebuild from scratch and compare to published | Released ≥ 2025 · ≤ 2.5 Å · clashscore < 5 · Rama outliers ≤ 0.1 % · rotamer outliers ≤ 1 % · 60–250 kDa. Excluded amyloids, whole capsids and single-chain spike deposits; one sequence per entry. | 198 | 9HJC, 11NX, 9DTU, 9EFZ, 9ZZI |

**Supplementary Table S2. Cryo-EM entries and deposited validation**

Entry metadata and deposited validation are from RCSB and wwPDB. These values may differ slightly from validation run within the exercise.

| PDB | EMDB | Structure | Released | Res. (Å) | kDa | Content | Deposited validation<br>clashscore / Rama outliers (%) / rotamer outliers (%) |
|---|---|---|---|---|---|---|---|
| **Batch 1 · user list (re-refinement)** | | | | | | | |
| 8YR2 | EMD-39533 | Structure of NET-Nisoxetine in outward-open state | 2024-05-29 | 2.89 | 151 | 2 chains · 41U, PIO, CLR, A1LX7 · ions: NA, CL | 21.87 / 0 / 5.36 |
| 8Y6K | EMD-38989 | Cryo-EM structure of full-length MICAL1 in the autoinhibited state | 2024-08-28 | 3.94 | 119 | FAD · ions: ZN | 21.64 / 0 / 0.33 |
| 9F6Z | EMD-17349 | Human N-deacetylase/N-sulfotransferase 1 homodimer | 2025-04-16 | 4.5 | 196 | 2 chains | 86.05 / 0.25 / 6.79 |
| 9BW9 | EMD-44962 | Tetrameric Complex of full-length HIV-1 integrase protein bound to the integrase binding domain of LEDGF/p75 | 2025-06-25 | 4.1 | 184 | 8 chains | 9.8 / 0 / 1.44 |
| 9JA6 | EMD-61291 | Cryo-EM structure of Tdk1 tetramer complex | 2024-11-13 | 4.4 | 154 | 4 chains | 19.69 / 0 / 9.56 |
| **Batch 2 · poor, 2025+, nucleic acid / protein–protein / drug** | | | | | | | |
| 9OWS | EMD-70942 | Structure of Geobacillus stearothermophilus RNase P holoenzyme in 5 mM Mg2+ | 2026-06-10 | 2.87 | 149 | RNA · 2 chains · ions: MG | 29.06 / 0 / 7.07 |
| 8YYV | EMD-39680 | A dimeric STAT1-DNA complex | 2025-03-26 | 3.07 | 102 | DNA · 3 chains | 27.79 / 6.96 / 11.26 |
| 9NRJ | EMD-49735 | abTCR bound to SAR444200, a novel anti-GPC3 T-cell engager, for the treatment of GPC3+ solid tumors | 2025-10-15 | 3.4 | 77 | 4 chains · NAG, FUC, BMA | 42.86 / 5.42 / 16.58 |

| PDB | EMDB | Structure | Released | Res. (Å) | kDa | Content | Deposited validation clashscore / Rama outliers (%) / rotamer outliers (%) |
|---|---|---|---|---|---|---|---|
| 9D0X | EMD-46465 | Cryo-EM structure of CDK2/CyclinE1 in complex with CRBN/DDB1 and Cpd 4 (local mask) | 2025-04-16 | 2.84 | 115 | 3 chains · A1A1I · ions: ZN | 31.49 / 0.79 / 5.38 |
| 9C0E | EMD-45081 | Phosphorylated human NKCC1_K289NA492E in complex with furosemide | 2025-02-12 | 2.7 | 266 | 2 chains · ATP, FUN · ions: K, CL, NA, MG | 21.19 / 0 / 4.51 |
| **Batch 3 · moderate quality, 2025+, 150–350 kDa** | | | | | | | |
| 8Z4O | EMD-39768 | MERS-CoV post-fusion S ectodomain trimer | 2025-10-22 | 3.79 | 180 | 3 chains · NAG | 9.66 / 2.37 / 4.37 |
| 9SKT | EMD-54974 | Cryo-EM structure of H. neapolitanus CsoSCA in oxidizing conditions, dimer, minor state | 2026-04-22 | 2.45 | 204 | 2 chains · ions: ZN | 8.52 / 0.96 / 4.83 |
| 9W0X | EMD-65520 | Cryo-EM structure of a Fungal XPR1 | 2026-03-18 | 2.9 | 181 | 2 chains · POV, 8PE, PO4 | 7.63 / 0.85 / 4.72 |
| 9DVT | EMD-47222 | Cryo-EM structure of RpaA bound to PkaiBC DNA and the CTD of the alpha subunit of RNAP from the cyanobacterium Synechococcus elongatus | 2025-10-15 | 3.74 | 202 | DNA · 6 chains | 8.91 / 1.01 / 3.57 |
| 8YS6 | EMD-39557 | Helicobacter pylori OorDABC in complex with Napabucasin | 2025-07-02 | 3.03 | 213 | 8 chains · SF4, A1D65, TPP · ions: MG | 6.93 / 0.65 / 3.95 |
| **Batch 4 · high quality, rebuilt de novo** | | | | | | | |
| 9HJC | EMD-52212 | Cryo-EM structure of thiamine-bound human TPK1 | 2025-03-12 | 2.1 | 63 | 2 chains · VIB · ions: MG | 2.21 / 0 / 0 |

| PDB | EMDB | Structure | Released | Res. (Å) | kDa | Content | Deposited validation clashscore / Rama outliers (%) / rotamer outliers (%) |
|---|---|---|---|---|---|---|---|
| 11NX | EMD-75874 | Rabbit muscle Aldolase (D2 symmetry) determined using the TFS Vitrobot Mark IV in the presence of 1x SurfACT | 2026-04-08 | 1.98 | 149 | 4 chains | 1.28 / 0 / 0.54 |
| 9DTU | EMD-47166 | Structure of the Measles virus Fusion protein ectodomain E170G E455G with mutated furin cleavage site in the pre-fusion conformation | 2025-10-22 | 2.05 | 174 | 3 chains · NAG | 2.6 / 0 / 0.44 |
| 9EFZ | EMD-47982 | Chicken BEST1 bound to GABA in an open state | 2025-04-09 | 1.95 | 205 | 5 chains · ABU · ions: CA, CL | 3.11 / 0 / 0.65 |
| 9ZZI | EMD-74998 | Cryo-EM structure of F-actin in the ADP state | 2026-06-24 | 2.06 | 212 | 5 chains · ADP · ions: MG | 1.96 / 0 / 0 |

**Supplementary Table S3. Software used in the exercise**

| Tool | Version | Used for |
|---|---|---|
| Phenix | 2.0-5936 (noCUDA) | real_space_refine (refinement), validation_cryoem, molprobity, cablam, reduce2 (riding H, NQH flips), elbow (ligand restraints), douse (waters), and the mmtbx API (per-residue CC, coordinate handling) |
| Coot | 0.9.8.95 (CCP4-9) | Scripted, no-graphics rotamer refit of remaining rotamer outliers (auto_fit_best_rotamer) |
| ModelAngelo | 1.0.18 | De novo building from map + sequence (batch 4). Upgraded from 1.0.12 into a new conda env model_angelo_1018 (Python 3.11, PyTorch 2.9.1 + CUDA 12.6 for driver 550); the old env is untouched. |
| ChimeraX + ISOLDE | 1.8 / 1.8 | Tested but not used: ISOLDE requires the GUI and no virtual display (Xvfb) is installed. |
| Q-score | Phenix 2.0 cctbx.maptbx.qscore | Q-score (Pintilie et al. 2020): 32 radial probes, shells 0.1–2.0 Å, run on the deposited primary map for every model |

| Tool | Version | Used for |
|---|---|---|
| Python (numpy, scipy) | Phenix-bundled | Custom scripts: trimming, false-connection fixing, water filtering, coordinate RMSD, model comparison, selection, tables |
| Data sources | — | Models from RCSB; maps and half-maps from the PDBj EMDB mirror (about 40× faster than EBI here); metadata from the RCSB Search and GraphQL APIs |

**Supplementary Table S4. Cryo-EM candidate selection rules**

| Rule | What it does |
|---|---|
| 1 · Geometry gate | A candidate must have a MolProbity score and clashscore no worse than the deposited model, \|Rama-Z\| ≤ 2 (Phenix's "good" range, which guards against over-restrained backbones), and zero broken peptide bonds (C–N ≤ 1.5 Å). |
| 2 · Keep the map fit | Among candidates that pass the gate, prefer those with CC_mask ≥ deposited − 0.01. Then keep only those within 0.02 of the best qualifying CC_mask, so a large map-fit gain is never traded for a small geometry gain. |
| 3 · Then best geometry | Take the lowest MolProbity score, breaking ties by higher CC_mask. If nothing passes the CC tolerance, take the lowest MolProbity score among gated candidates within 0.01 of the best CC_mask. |
| Trimming (CC < 0.2) | A residue is flagged when its whole-residue CC is below 0.2. Runs of 2 or more flagged residues, and flagged chain termini, are deleted. An isolated flagged residue keeps its backbone (so the chain doesn't break) and is cut back to Cβ; isolated Gly/Ala are kept and flagged. Nucleotides follow the same rule but are never truncated. Ligands, ions, water and glycans are never trimmed. |
| Coot guard | Coot's rotamer fit can shift the backbone. After Coot broke peptide bonds in 8Y6K and 9BW9, only its side-chain conformations were kept, re-superposed onto the original N/Cα/C atoms. |
| Map weight | The batch-1 scan showed that higher map weights always raised CC at a large geometry cost (9JA6 at w = 8 had clashscore 106). This is overfitting, so automatic weighting was used from then on. |
| B-factors | When CC dropped with coordinates unchanged, the cause was B-factor refinement. Both options were then run for every entry and the selection rule chose between them. |

### Cryo-EM re-refinement:

#### Supplementary Table S5. Batch 1 geometry and map-fit metrics

Each pair is deposited → refined. The symbols ▲ and ▼ reproduce the original report. Rama-Z is judged by distance from zero.

| Metric | 8YR2 · 2.89 Å | 8Y6K · 3.94 Å | 9F6Z · 4.5 Å | 9BW9 · 4.1 Å | 9JA6 · 4.4 Å |
|---|---|---|---|---|---|
| Protein residues | 1130 → 1130 | 741 → 729 | 1609 → 1560 | 1282 → 1282 | 588 → 588 |
| Bond RMSD (Å) | 0.008 → 0.005 | 0.007 → 0.004 | 0.018 → 0.006 | 0.004 → 0.004 | 0.008 → 0.004 |
| Angle RMSD (°) | 1.31 → 1.06 | 1.22 → 0.96 | 2.14 → 1.01 | 0.94 → 0.95 | 1.16 → 0.93 |
| MolProbity score | 2.66 → 1.82 ▲ | 2.16 → 1.76 ▲ | 3.03 → 2.14 ▲ | 1.96 → 1.65 ▲ | 2.67 → 1.70 ▲ |
| Clashscore | 21.6 → 6.8 ▲ | 21.5 → 7.2 ▲ | 86.0 → 16.1 ▲ | 9.6 → 5.5 ▲ | 19.6 → 7.3 ▲ |
| Rotamer outliers (%) | 5.36 → 1.68 ▲ | 0.33 → 1.18 ▼ | 6.79 → 0.00 ▲ | 1.44 → 0.00 ▲ | 9.56 → 0.00 ▲ |
| Rama favored (%) | 95.9 → 96.0 ▲ | 95.2 → 95.5 ▲ | 98.8 → 93.5 ▼ | 95.2 → 94.9 ▼ | 97.2 → 95.7 ▼ |
| Rama outliers (%) | 0.00 → 0.00 | 0.00 → 0.00 | 0.25 → 0.00 ▲ | 0.00 → 0.00 | 0.00 → 0.00 |
| Rama-Z | -1.38 → 1.28 ▲ | -0.44 → -0.27 ▲ | -0.08 → -0.69 ▼ | -0.88 → 0.16 ▲ | -1.98 → 1.69 ▲ |
| CaBLAM outliers (%) | 1.2 → 1.4 ▼ | 1.4 → 1.4 = | 1.8 → 2.3 ▼ | 3.0 → 2.5 ▲ | 0.0 → 0.7 ▼ |
| Cβ deviations | 0 → 0 | 0 → 0 = | 4 → 0 ▲ | 0 → 0 = | 0 → 0 |
| CC_mask | 0.628 → 0.834 ▲ | 0.590 → 0.582 | 0.397 → 0.624 ▲ | 0.746 → 0.754 | 0.597 → 0.766 ▲ |
| CC_box | 0.489 → 0.625 ▲ | 0.657 → 0.652 | 0.651 → 0.746 ▲ | 0.676 → 0.681 | 0.688 → 0.711 ▲ |
| CC main chain | 0.643 → 0.827 ▲ | 0.591 → 0.587 | 0.454 → 0.650 ▲ | 0.732 → 0.740 | 0.624 → 0.713 ▲ |
| CC side chain | 0.567 → 0.771 ▲ | 0.597 → 0.592 | 0.505 → 0.671 ▲ | 0.720 → 0.728 | 0.630 → 0.702 ▲ |
| Residues CC < 0.2 | 2 → 0 ▲ | 14 → 5 ▲ | 204 → 37 ▲ | 0 → 0 | 0 → 0 |
| Broken peptides | 0 → 0 | 0 → 0 | 0 → 0 | 0 → 0 | 0 → 0 |
| Cα RMSD vs deposited (Å) | 0.35 | 0.18 | 0.66 | 0.23 | 0.56 |

| Metric | 8YR2 · 2.89 Å | 8Y6K · 3.94 Å | 9F6Z · 4.5 Å | 9BW9 · 4.1 Å | 9JA6 · 4.4 Å |
|---|---|---|---|---|---|
| Max Cα shift (Å) | 1.27 | 1.17 | 1.90 | 0.92 | 1.31 |
| Trim: deleted / to Cβ / flagged | 0 / 0 / 0 | 12 / 4 / 4 | 49 / 37 / 10 | 0 / 0 / 0 | 0 / 0 / 0 |

**Supplementary Table S6. Batch 2 geometry and map-fit metrics**

Each pair is deposited → refined. The symbols ▲ and ▼ reproduce the original report. Rama-Z is judged by distance from zero.

| Metric | 9OWS · 2.87 Å | 8YYV · 3.07 Å | 9NRJ · 3.4 Å | 9D0X · 2.84 Å | 9C0E · 2.7 Å |
|---|---|---|---|---|---|
| Protein residues | 116 → 116 | 1106 → 1106 | 672 → 672 | 896 → 896 | 1818 → 1818 |
| Bond RMSD (Å) | 0.011 → 0.006 | 0.014 → 0.005 | 0.051 → 0.005 | 0.009 → 0.006 | 0.010 → 0.004 |
| Angle RMSD (°) | 1.82 → 0.67 | 2.78 → 0.94 | 7.28 → 1.09 | 1.46 → 1.09 | 1.02 → 0.97 |
| MolProbity score | 3.04 → 2.06 ▲ | 3.52 → 1.75 ▲ | 3.73 → 2.26 ▲ | 3.02 → 2.01 ▲ | 2.62 → 1.34 ▲ |
| Clashscore | 29.0 → 9.3 ▲ | 27.79 → 2.6 ▲ | 42.9 → 6.1 ▲ | 31.6 → 6.0 ▲ | 21.0 → 2.1 ▲ |
| Rotamer outliers (%) | 7.07 → 0.00 ▲ | 11.26 → 1.48 ▲ | 16.58 → 3.45 ▲ | 5.38 → 1.88 ▲ | 4.51 → 1.09 ▲ |
| Rama favored (%) | 93.0 → 89.5 ▼ | 77.4 → 88.9 ▲ | 83.4 → 91.0 ▲ | 92.1 → 92.3 ▲ | 95.6 → 95.3 ▼ |
| Rama outliers (%) | 0.00 → 0.00 | 6.99 → 1.29 ▲ | 5.42 → 0.45 ▲ | 0.68 → 0.34 ▲ | 0.00 → 0.00 |
| Rama-Z | -0.85 → -0.34 ▲ | -6.21 → -1.84 ▲ | -4.68 → -2.64 ▲ | -1.28 → -0.57 ▲ | 2.11 → 1.28 ▲ |
| CaBLAM outliers (%) | 2.7 → 2.7 | 8.9 → 5.9 ▲ | 6.2 → 4.7 ▲ | 4.4 → 3.6 ▲ | 3.2 → 3.3 ▼ |
| Cβ deviations | 1 → 0 ▲ | 80 → 0 ▲ | 384 → 0 ▲ | 1 → 0 ▲ | 0 → 0 |
| CC_mask | 0.806 → 0.878 ▲ | 0.816 → 0.811 | 0.792 → 0.783 ▼ | 0.841 → 0.842 | 0.898 → 0.899 |
| CC_box | 0.889 → 0.913 ▲ | 0.506 → 0.501 | 0.671 → 0.673 | 0.806 → 0.807 | 0.734 → 0.735 |
| CC main chain | 0.788 → 0.841 ▲ | 0.844 → 0.836 | 0.771 → 0.748 ▼ | 0.841 → 0.843 | 0.882 → 0.883 |
| CC side chain | 0.821 → 0.880 ▲ | 0.783 → 0.776 | 0.735 → 0.722 ▼ | 0.833 → 0.834 | 0.874 → 0.872 |

| Metric | 9OWS · 2.87 Å | 8YYV · 3.07 Å | 9NRJ · 3.4 Å | 9D0X · 2.84 Å | 9C0E · 2.7 Å |
|---|---|---|---|---|---|
| Residues CC < 0.2 | 1 → 1 | 0 → 0 | 0 → 1 ▼ | 0 → 0 = | 0 → 0 = |
| Broken peptides | 0 → 0 | 0 → 0 | 3 → 0 ▲ | 0 → 0 | 0 → 0 |
| Cα RMSD vs deposited (Å) | 0.54 | 0.24 | 0.44 | 0.17 | 0.10 |
| Max Cα shift (Å) | 1.02 | 0.73 | 1.67 | 0.72 | 0.48 |
| Trim: deleted / to Cβ / flagged | 0 / 0 / 1 | 0 / 0 / 0 | 0 / 0 / 1 | 0 / 0 / 0 | 0 / 0 / 0 |

**Supplementary Table S7. Batch 3 geometry and map-fit metrics**

Each pair is deposited → refined. The symbols ▲ and ▼ reproduce the original report. Rama-Z is judged by distance from zero.

| Metric | 8Z4O · 3.79 Å | 9SKT · 2.45 Å | 9W0X · 2.9 Å | 9DVT · 3.74 Å | 8YS6 · 3.03 Å |
|---|---|---|---|---|---|
| Protein residues | 1073 → 1073 | 942 → 942 | 1190 → 1190 | 607 → 607 | 1874 → 1874 |
| Bond RMSD (Å) | 0.014 → 0.005 | 0.014 → 0.004 | 0.012 → 0.004 | 0.003 → 0.004 | 0.024 → 0.004 |
| Angle RMSD (°) | 2.32 → 1.00 | 2.13 → 0.95 | 1.36 → 0.91 | 0.66 → 0.92 | 1.53 → 1.08 |
| MolProbity score | 2.37 → 2.08 ▲ | 2.47 → 0.84 ▲ | 2.32 → 1.71 ▲ | 2.13 → 1.78 ▲ | 2.33 → 1.43 ▲ |
| Clashscore | 9.6 → 4.7 ▲ | 8.4 → 0.9 ▲ | 7.6 → 2.1 ▲ | 8.8 → 6.7 ▲ | 6.9 → 2.0 ▲ |
| Rotamer outliers (%) | 4.37 → 2.89 ▲ | 4.83 → 0.51 ▲ | 4.72 → 2.69 ▲ | 3.57 → 1.50 ▲ | 3.95 → 0.83 ▲ |
| Rama favored (%) | 93.0 → 91.9 ▼ | 91.8 → 97.8 ▲ | 94.2 → 94.0 ▼ | 96.4 → 95.9 ▼ | 91.8 → 92.8 ▲ |
| Rama outliers (%) | 2.37 → 0.38 ▲ | 0.96 → 0.00 ▲ | 0.85 → 0.34 ▲ | 1.02 → 0.17 ▲ | 0.65 → 0.16 ▲ |
| Rama-Z | 1.42 → -0.10 ▲ | -3.55 → 1.73 ▲ | 0.82 → 1.56 ▼ | -0.56 → 0.73 ▼ | -3.81 → -0.83 ▲ |
| CaBLAM outliers (%) | 4.3 → 4.6 ▼ | 2.4 → 0.9 ▲ | 3.6 → 3.1 ▲ | 1.0 → 2.2 ▼ | 3.3 → 3.7 ▼ |
| Cβ deviations | 30 → 0 ▲ | 16 → 0 ▲ | 8 → 0 ▲ | 0 → 0 | 1 → 0 ▲ |
| CC_mask | 0.793 → 0.790 | 0.841 → 0.885 ▲ | 0.854 → 0.854 | 0.708 → 0.745 ▲ | 0.852 → 0.855 |

| Metric | 8Z4O · 3.79 Å | 9SKT · 2.45 Å | 9W0X · 2.9 Å | 9DVT · 3.74 Å | 8YS6 · 3.03 Å |
|---|---|---|---|---|---|
| CC_box | 0.783 → 0.782 | 0.597 → 0.688 ▲ | 0.686 → 0.686 | 0.736 → 0.761 ▲ | 0.750 → 0.754 |
| CC main chain | 0.793 → 0.791 | 0.822 → 0.871 ▲ | 0.869 → 0.869 | 0.673 → 0.731 ▲ | 0.845 → 0.849 |
| CC side chain | 0.771 → 0.772 | 0.770 → 0.832 ▲ | 0.835 → 0.835 | 0.702 → 0.737 ▲ | 0.816 → 0.823 |
| Residues CC < 0.2 | 0 → 0 | 3 → 0 ▲ | 0 → 0 | 2 → 0 ▲ | 1 → 4 ▼ |
| Broken peptides | 0 → 0 | 0 → 0 | 0 → 0 | 0 → 0 | 0 → 0 |
| Cα RMSD vs deposited (Å) | 0.34 | 0.19 | 0.16 | 0.38 | 0.24 |
| Max Cα shift (Å) | 1.20 | 0.62 | 0.71 | 1.34 | 0.99 |
| Trim: deleted / to Cβ / flagged | 0 / 0 / 0 | 0 / 0 / 0 | 0 / 0 / 0 | 0 / 0 / 1 | 0 / 0 / 0 |

**Supplementary Table S8. Q-scores for re-refined cryo-EM entries**

Before and after values use the same Phenix implementation and map. The wwPDB value is shown for reference, not as a directly interchangeable score.

| PDB | Res. (Å) | wwPDB Q (deposited) | Q all: deposited → refined | Q backbone | Q side chain |
|---|---|---|---|---|---|
| **Batch 1 · user list (re-refinement)** | | | | | |
| 8YR2 | 2.89 | 0.470 | 0.561 → 0.628 ▲ | 0.571 → 0.622 ▲ | 0.540 → 0.644 ▲ |
| 8Y6K | 3.94 | 0.183 | 0.221 → 0.231 ▲ | 0.282 → 0.291 ▲ | 0.152 → 0.165 ▲ |
| 9F6Z | 4.5 | 0.097 | 0.119 → 0.227 ▲ | 0.165 → 0.249 ▲ | 0.075 → 0.206 ▲ |
| 9BW9 | 4.1 | 0.332 | 0.409 → 0.422 ▲ | 0.412 → 0.416 | 0.406 → 0.427 ▲ |
| 9JA6 | 4.4 | 0.142 | 0.173 → 0.214 ▲ | 0.345 → 0.396 ▲ | 0.013 → 0.045 ▲ |
| **Batch 2 · poor, 2025+, nucleic acid / protein–protein / drug** | | | | | |
| 9OWS | 2.87 | 0.394 | 0.477 → 0.497 ▲ | 0.541 → 0.598 ▲ | 0.508 → 0.610 ▲ |
| 8YYV | 3.07 | 0.548 | 0.653 → 0.652 | 0.644 → 0.631 ▼ | 0.659 → 0.661 ▲ |

| PDB | Res. (Å) | wwPDB Q (deposited) | Q all: deposited → refined | Q backbone | Q side chain |
|---|---|---|---|---|---|
| 9NRJ | 3.4 | 0.441 | 0.531 → 0.568 ▲ | 0.599 → 0.583 ▼ | 0.467 → 0.561 ▲ |
| 9D0X | 2.84 | 0.536 | 0.621 → 0.621 | 0.639 → 0.637 | 0.602 → 0.605 |
| 9C0E | 2.7 | 0.600 | 0.711 → 0.712 | 0.694 → 0.695 | 0.732 → 0.732 |
| **Batch 3 · moderate quality, 2025+, 150–350 kDa** | | | | | |
| 8Z4O | 3.79 | 0.426 | 0.518 → 0.510 | 0.540 → 0.514 ▼ | 0.492 → 0.508 ▲ |
| 9SKT | 2.45 | 0.654 | 0.763 → 0.760 | 0.769 → 0.759 ▼ | 0.758 → 0.761 |
| 9W0X | 2.9 | 0.563 | 0.668 → 0.666 | 0.639 → 0.639 | 0.688 → 0.686 |
| 9DVT | 3.74 | 0.317 | 0.389 → 0.453 ▲ | 0.444 → 0.475 ▲ | 0.339 → 0.461 ▲ |
| 8YS6 | 3.03 | 0.531 | 0.632 → 0.648 ▲ | 0.627 → 0.626 | 0.639 → 0.673 ▲ |

### De novo model building

**Supplementary Table S9. Q-scores for de novo models and published models**

Each pair is de novo / published and is scored with the same implementation and map.

| PDB | Res. (Å) | wwPDB Q (published) | Q all: de novo / published | Q backbone | Q side chain |
|---|---|---|---|---|---|
| 9HJC | 2.1 | 0.697 | 0.797 / 0.796 | 0.798 / 0.802 ▼ | 0.797 / 0.790 |
| 11NX | 1.98 | 0.747 | 0.866 / 0.839 ▲ | 0.876 / 0.863 ▲ | 0.856 / 0.811 ▲ |
| 9DTU | 2.05 | 0.724 | 0.818 / 0.818 | 0.831 / 0.838 | 0.805 / 0.804 |
| 9EFZ | 1.95 | 0.750 | 0.862 / 0.854 ▲ | 0.867 / 0.863 ▲ | 0.857 / 0.847 ▲ |
| 9ZZI | 2.06 | 0.738 | 0.863 / 0.839 ▲ | 0.871 / 0.857 ▲ | 0.856 / 0.820 ▲ |

### Half-map validation:

**Supplementary Table S10. Half-map cross-validation results**

Work and free scores were calculated by refining against one deposited half-map and scoring against the other. The two directions were averaged.

| PDB | Res. (Å) | CC_mask work / free | CC gap | FSCavg work / free | FSC gap | Final model vs half 1 / half 2 |
|---|---|---|---|---|---|---|
| **Batch 1 · user list (re-refinement)** | | | | | | |
| 8YR2 | 2.89 | 0.816 / 0.813 | +0.003 | 0.541 / 0.529 | +0.012 | 0.752 / 0.754 |
| 8Y6K | 3.94 | 0.675 / 0.663 | +0.011 | 0.400 / 0.368 | +0.032 | 0.598 / 0.595 |
| 9F6Z | 4.5 | 0.688 / 0.683 | +0.005 | 0.285 / 0.265 | +0.020 | 0.631 / 0.626 |
| 9BW9 | 4.1 | 0.660 / 0.653 | +0.008 | 0.561 / 0.543 | +0.018 | 0.654 / 0.660 |
| 9JA6 | 4.4 | 0.828 / 0.814 | +0.014 | 0.403 / 0.373 | +0.030 | 0.795 / 0.803 |
| **Batch 2 · poor, 2025+, nucleic acid / protein–protein / drug** | | | | | | |
| 9OWS | 2.87 | 0.842 / 0.838 | +0.004 | 0.697 / 0.683 | +0.013 | 0.858 / 0.858 |
| 8YYV | 3.07 | 0.867 / 0.861 | +0.006 | 0.602 / 0.588 | +0.013 | 0.875 / 0.874 |
| 9NRJ | 3.4 | 0.761 / 0.755 | +0.006 | 0.624 / 0.606 | +0.018 | 0.677 / 0.675 |
| 9D0X | 2.84 | 0.794 / 0.779 | +0.015 | 0.567 / 0.528 | +0.039 | 0.801 / 0.801 |
| 9C0E | 2.7 | 0.826 / 0.820 | +0.005 | 0.619 / 0.599 | +0.020 | 0.835 / 0.834 |
| **Batch 3 · moderate quality, 2025+, 150–350 kDa** | | | | | | |
| 8Z4O | 3.79 | 0.719 / 0.702 | +0.017 | 0.607 / 0.572 | +0.036 | 0.714 / 0.722 |
| 9SKT | 2.45 | 0.693 / 0.692 | +0.001 | 0.579 / 0.565 | +0.014 | 0.703 / 0.702 |
| 9W0X | 2.9 | 0.806 / 0.801 | +0.006 | 0.635 / 0.620 | +0.015 | 0.807 / 0.806 |
| 9DVT | 3.74 | 0.615 / 0.593 | +0.022 | 0.555 / 0.511 | +0.044 | 0.612 / 0.613 |

| PDB | Res. (Å) | CC_mask work / free | CC gap | FSCavg work / free | FSC gap | Final model vs half 1 / half 2 |
|---|---|---|---|---|---|---|
| 8YS6 | 3.03 | 0.761 / 0.753 | +0.008 | 0.643 / 0.622 | +0.021 | 0.764 / 0.765 |
| **Batch 4 · high quality, rebuilt de novo** | | | | | | |
| 9HJC | 2.1 | 0.843 / 0.841 | +0.001 | 0.508 / 0.493 | +0.015 | 0.825 / 0.827 |
| 11NX | 1.98 | 0.905 / 0.904 | +0.002 | 0.674 / 0.660 | +0.014 | 0.887 / 0.887 |
| 9DTU | 2.05 | 0.907 / 0.906 | +0.001 | 0.606 / 0.597 | +0.009 | 0.887 / 0.887 |
| 9EFZ | 1.95 | 0.888 / 0.883 | +0.005 | 0.639 / 0.614 | +0.025 | 0.895 / 0.894 |
| 9ZZI | 2.06 | 0.872 / 0.871 | +0.001 | 0.567 / 0.561 | +0.006 | 0.818 / 0.818 |

### De novo model building

#### Supplementary Table S11. De novo versus published model metrics

Each pair is de novo / published. For 9ZZI, metrics use the five subunits matching the published model. The de novo exercise tested protein building from maps and sequences. Ligands, ions and glycans were outside its scope and absent from the de novo candidates. Thus, these protein-only metrics do not establish complete chemical models. Deposited nonprotein components in 9DTU, 9HJC, 9EFZ and 9ZZI are described in the main text.

| De novo / published | 9HJC | 11NX | 9DTU | 9EFZ | 9ZZI |
|---|---|---|---|---|---|
| Resolution (Å) | 2.1 | 1.98 | 2.05 | 1.95 | 2.06 |
| Protein residues | 465 / 486 | 1368 / 1372 | 1346 / 1350 | 1704 / 1700 | 1846 / 1855 |
| Waters | 273 / 0 | 976 / 1794 | 897 / 400 | 925 / 382 | 1325 / 205 |
| MolProbity score | 1.38 / 1.10 ▼ | 1.15 / 0.95 ▼ | 0.75 / 1.05 ▲ | 1.17 / 1.10 ▼ | 1.00 / 1.05 ▲ |
| Clashscore | 2.3 / 2.2 ▼ | 2.0 / 1.2 ▼ | 0.8 / 2.6 ▲ | 3.0 / 3.1 ▲ | 2.2 / 1.9 ▼ |
| Rotamer outliers (%) | 0.98 / 0.00 ▼ | 1.26 / 0.54 ▼ | 0.53 / 0.44 ▼ | 0.26 / 0.65 ▲ | 0.32 / 0.00 ▼ |
| Rama favored (%) | 94.7 / 97.5 ▼ | 97.5 / 97.4 ▲ | 98.0 / 99.3 ▼ | 97.6 / 99.1 ▼ | 98.0 / 97.5 ▲ |

| De novo / published | 9HJC | 11NX | 9DTU | 9EFZ | 9ZZI |
|---|---|---|---|---|---|
| Rama-Z | 0.16 / 1.01 ▲ | 0.89 / 0.83 ▼ | 1.08 / 0.94 ▼ | 0.45 / 0.35 ▼ | 1.21 / -1.74 ▲ |
| CaBLAM outliers (%) | 2.3 / 2.7 ▲ | 1.6 / 1.4 ▼ | 1.2 / 0.5 ▼ | 0.1 / 0.3 ▲ | 0.4 / 0.3 ▼ |
| CC_mask | 0.793 / 0.802 ▼ | 0.875 / 0.852 ▲ | 0.845 / 0.852 ▼ | 0.893 / 0.903 ▼ | 0.927 / 0.903 ▲ |
| CC main chain | 0.776 / 0.767 ▲ | 0.878 / 0.865 ▲ | 0.840 / 0.839 ▲ | 0.893 / 0.898 ▼ | 0.922 / 0.890 ▲ |
| Raw ModelAngelo: residues · clashscore | 469 · 77.5 | 1368 · 48.5 | 1352 · 37.0 | 1704 · 52.1 | 3207 · 47.5 |
| Published residues recovered | 95.5 % | 99.4 % | 98.8 % | 100 % | 99.2 % |
| Same residue type at matched positions | 100 % | 100 % | 100 % | 100 % | 100 % |
| Cα RMSD, matched (Å) | 0.16 | 0.15 | 0.21 | 0.10 | 0.19 |
| All-atom RMSD, same residues (Å) | 0.53 | 0.66 | 0.63 | 0.37 | 0.51 |
| Published waters recovered (≤ 1 Å) | – | 47 % | 86.5 % | 87.2 % | 78.5 % |
| Our waters in published ligand/ion sites | 3 | 0 | 8 | 17 | 0 |

### In-house structures

**Supplementary Table S12. In-house entry metadata**

| PDB | Method | Structure | Res. (Å) | EMDB / space group · cell | Deposited (wwPDB report) |
|---|---|---|---|---|---|
| 7YQM | cryo-EM | 2.9-angstrom cryo-EM structure of Ecoli malate synthase G | 2.89 | EMD-34029 | clash 1.78 · Q 0.548 |
| 7YQN | X-ray | Crystal structure of Ecoli malate synthase G | 1.6 | P 21 21 21 · 98.9 / 101.2 / 170.2 Å | R 0.156 / 0.172 · RSRZ 2.4 % |
| 9LQM | X-ray | Crystal structure of synthetic ubiquitin variant R4 designed by ProteinMPNN | 1.39 | P 32 2 1 · 50.8 / 50.8 / 102.7 Å | R 0.164 / 0.196 · RSRZ 5.6 % |
| 9LQK | X-ray | Crystal structure of synthetic ubiquitin variant R10 designed by ProteinMPNN | 1.55 | I 2 2 2 · 57.6 / 62.5 / 89.8 Å | R 0.191 / 0.229 · RSRZ 9.5 % |

**Supplementary Table S13. Cryo-EM re-refinement of 7YQM**

The deposited 7YQM model retained better geometry and was not superseded by the exercise candidate.

| Metric (deposited → refined) | 7YQM · 2.89 Å |
| --- | --- |
| MolProbity score | 1.25 → 1.33 ▼ |
| Clashscore | 1.69 → 2.41 ▼ |
| Rotamer outliers (%) | 0.17 → 0.50 ▼ |
| Rama favored (%) | 95.28 → 95.53 ▲ |
| CaBLAM outliers (%) | 1.1 → 1.0 ▲ |
| CC_mask | 0.780 → 0.786 |
| CC side chain | 0.731 → 0.740 ▲ |
| Residues CC < 0.2 | 7 → 3 ▲ |
| Q-score all / backbone / side chain | 0.649 / 0.636 / 0.663 → 0.663 / 0.640 / 0.686 |
| Half-map test: CC gap / FSCavg gap | 0.019 / 0.030 |
| Cα RMSD vs deposited / max shift (Å) | 0.12 / 0.38 |
| Trim: deleted / to Cβ | 3 / 2 |

### Crystal structure workflow

#### Supplementary Table S14. Crystal structure decision rules

| Rule | What it does |
|---|---|
| RSRZ (wwPDB style) | Per-residue RSR and RSCC from locally computed 2mFo−DFc and Fc maps (EDS radius). The reference mean/sd of RSR per residue type and resolution were recovered exactly from each entry's official wwPDB validation report, and the local RSR was calibrated to the wwPDB RSR on the published model (reduced-major-axis fit; r = 0.73–0.88). For the published models the local estimate reproduces the official value within 0.9 points for 7YQN and 9LQM, and reads 2.5 points low for 9LQK. The report is used only for this reference, never as a model. |
| Omit and rebuild | Segments with RSCC far below their neighbours were deleted (ubiquitin C-terminal tail 73–76, still in the wild-type path; MSG loop 151–156 and residues 3–4), waters near them removed, and 2mFo−DFc maps recomputed after refinement without them. Loops were rebuilt with phenix.fit_loops, which works headless; Coot's fit_gap fails without a GUI in this version. |
| Grow termini one residue per round | Coot adds the next sequence residue at each open end, then riding H and 3 macro-cycles of refinement. The residue stays only if RSCC ≥ 0.8 and RSRZ ≤ 3; otherwise it is removed and that end stops. Adding all residues at once and trimming afterwards failed: outer residues were placed before the map could show them. |
| Water search | phenix.refine's default only accepts waters within 3.2 Å of backbone atoms, which misses waters bound to side chains or to other waters. Any non-H atom is now a valid partner (2.4–3.4 Å), picking runs every macro-cycle, and the peak (mFo−DFc ≥ 3σ), 2mFo−DFc ≥ 1σ, map-CC ≥ 0.7 and B ≤ 80 Å² filters stay. Rounds continue while $R_{free}$ drops by ≥ 0.001. |
| ADP model | Isotropic, TLS (one group per chain) + isotropic, and anisotropic runs from the same start. The lowest $R_{free}$ wins, but a more complex scheme must beat every simpler one by ≥ 0.005. |
| Alternate conformations | phenix.create_alt_conf, then the alt-conf model and a control get identical refinement. Accept only if $R_{free}$ drops by ≥ 0.002, the $R_{free} - R_{work}$ gap grows by ≤ 0.01 and the clashscore is not worse. |
| Clash clean-up | Waters within 2.9 Å of a carbon or 2.1 Å of a hydrogen removed; rotamer outliers and residues in ≥ 3 clashes refitted in Coot (side chains only kept); final polish at a fixed tight geometry weight (wxc_scale 0.25). |

**Supplementary Table S15. Crystal first-pass final and published metrics**

Each value is first pass → final · published. The models were scored H-free on the same data and test set. RSRZ is a calibrated local estimate; official wwPDB values for published models are in brackets.

| Metric | 7YQN · 1.6 Å | 9LQM · 1.39 Å | 9LQK · 1.55 Å |
|---|---|---|---|
| $R_{work}$ | 0.172 → 0.151 ▲ · 0.156 | 0.202 → 0.193 ▲ · 0.168 | 0.229 → 0.205 ▲ · 0.192 |
| $R_{free}$ | 0.202 → 0.182 ▲ · 0.173 | 0.237 → 0.207 ▲ · 0.197 | 0.272 → 0.238 ▲ · 0.229 |
| MolProbity score | 2.19 → 1.29 ▲ · 0.87 | 1.73 → 0.84 ▲ · 0.74 | 1.77 → 1.43 ▲ · 1.01 |
| Clashscore | 20.84 → 5.44 ▲ · 1.37 | 5.89 → 1.20 ▲ · 0.76 | 7.10 → 7.84 ▼ · 2.29 |
| Rotamer outliers (%) | 2.64 → 0.00 ▲ · 0.08 | 0.69 → 0.00 ▲ · 0.68 | 2.08 → 0.72 ▲ · 0.00 |
| Rama favored (%) | 97.73 → 98.30 ▲ · 98.80 | 93.92 → 98.62 ▲ · 98.73 | 97.30 → 99.30 ▲ · 100.00 |
| CaBLAM outliers (%) | 0.8 → 0.4 ▲ · 0.5 | 2.1 → 0.0 ▲ · 0.0 | 1.4 → 0.0 ▲ · 0.0 |
| $R_{work}$ / $R_{free}$ with riding H (final) | 0.141 / 0.175 | 0.185 / 0.202 | 0.199 / 0.230 |
| RSRZ outliers (%) | 4.87 → 3.95 ▲ · 3.22 [2.38] | 7.89 → 3.36 ▲ · 5.59 [5.59] | 13.16 → 1.37 ▲ · 6.96 [9.49] |
| Mean RSCC (protein) | 0.933 → 0.947 ▲ · 0.950 | 0.915 → 0.946 ▲ · 0.955 | 0.871 → 0.915 ▲ · 0.918 |
| Protein residues | 1418 → 1416 · 1430 | 152 → 149 · 161 | 152 → 146 · 158 |
| Waters (final · published) | 2272 · 1881 | 247 · 237 | 190 · 175 |
| ADP model (final) | anisotropic (non-H, non-water) | TLS (one group per chain) + isotropic | TLS (one group per chain) + isotropic |

**Supplementary Table S16. Crystal second-pass refinement steps**

The R-work and R-free in this table are from phenix.refine with riding hydrogen atoms.

| Step | What was done | R-work / R-free |
|---|---|---|
| **7YQN · published R-free on the same data 0.173** | | |
| 1 | First pass (MR, rebuild, ligands, waters) | 0.1629 / 0.1945 |
| 2 | Loop 151–156 omitted + rebuilt; distorted GOL/BME removed | 0.1670 / 0.1982 |
| 3 | N-terminus regrow test (residue 4 rejected) | 0.1669 / 0.1987 |
| 4 | ADP model: anisotropic | 0.1343 / 0.1819 |
| 5 | Loop 300–310 regrow (V310 kept) | 0.1394 / 0.1845 |
| 6 | Waters: broadened search | 0.1333 / 0.1791 |
| 7 | Clash clean-up + final polish | 0.1407 / 0.1752 |
| **9LQM · published R-free on the same data 0.197** | | |
| 1 | First pass (MR, thread, refine, waters) | 0.1948 / 0.2315 |
| 2 | Termini omitted + regrown | 0.1873 / 0.2201 |
| 3 | Waters: broadened search | 0.1634 / 0.1972 |
| 4 | ADP model: TLS | 0.1777 / 0.1979 |
| 5 | Alt-conf test (rejected; control kept) | 0.1747 / 0.2002 |
| 6 | Clash clean-up + final polish | 0.1848 / 0.2018 |
| **9LQK · published R-free on the same data 0.229** | | |
| 1 | First pass (MR, thread, refine, waters) | 0.2219 / 0.2678 |
| 2 | Termini omitted + regrown | 0.2220 / 0.2569 |
| 3 | ADP model: TLS | 0.1997 / 0.2368 |

| Step | What was done | R-work / R-free |
|---|---|---|
| 4 | Waters: broadened search | 0.1913 / 0.2304 |
| 5 | Alt-conf test (rejected; control kept) | 0.1956 / 0.2288 |
| 6 | Clash clean-up + final polish | 0.1989 / 0.2301 |

**Supplementary Table S17. Crystal parameter tests and decisions**

| Test | Result | Decision |
|---|---|---|
| 7YQN ADP | iso $R_{work}$ 0.1593 $R_{free}$ 0.1909 gap 0.0316 \| tls $R_{work}$ 0.1568 $R_{free}$ 0.1892 gap 0.0324 \| aniso $R_{work}$ 0.1343 $R_{free}$ 0.1819 gap 0.0476 | anisotropic (non-H, non-water); chosen over isotropic / TLS by $R_{free}$ |
| 9LQM ADP | iso $R_{work}$ 0.1913 $R_{free}$ 0.2105 gap 0.0192 \| tls $R_{work}$ 0.1777 $R_{free}$ 0.1979 gap 0.0202 \| aniso $R_{work}$ 0.1600 $R_{f\backslash ree}$ 0.1969 gap 0.0369 | TLS (one group per chain) + isotropic; chosen over isotropic / anisotropic by $R_{free}$ |
| 9LQM alt conf | alt-conf $R_{work}$ 0.1776 $R_{free}$ 0.2021 gap 0.0245 clash 5.02 \| control $R_{work}$ 0.1747 $R_{free}$ 0.2002 gap 0.0255 clash 9.64 | Rejected: create_alt_conf gave a near-whole-model two-conformer ensemble (all residues split, over half by > 0.5 Å); $R_{free}$ rose. The published models have 0 (9LQM), 3 (9LQK) and 46 (7YQN) residues with alternates. |
| 9LQK ADP | iso $R_{work}$ 0.2171 $R_{free}$ 0.2571 gap 0.0400 \| tls $R_{work}$ 0.1997 $R_{free}$ 0.2368 gap 0.0371 \| aniso $R_{work}$ 0.1850 $R_{free}$ 0.2456 gap 0.0606 | TLS (one group per chain) + isotropic; chosen over isotropic / anisotropic by $R_{free}$ |
| 9LQK alt conf | alt-conf $R_{work}$ 0.1888 $R_{free}$ 0.2321 gap 0.0433 clash 11.56 \| control $R_{work}$ 0.1956 $R_{free}$ 0.2288 gap 0.0332 clash 16.09 | Rejected: create_alt_conf gave a near-whole-model two-conformer ensemble (all residues split, over half by > 0.5 Å); $R_{free}$ rose. The published models have 0 (9LQM), 3 (9LQK) and 46 (7YQN) residues with alternates. |

**Supplementary Table S18. Crystal model composition compared with published entries**

For 7YQN, the exercise final model contained two GOL, two BME and two NA, while the published entry contains four GOL, two BME, three CL and two NA. The automated workflow therefore did not recover every deposited small-molecule or ion site.

| Ours vs published | 7YQN | 9LQM | 9LQK |
|---|---|---|---|
| Published residues / AI-supervised | 1430 / 1416 | 161 / 149 | 158 / 146 |
| Published residues recovered | 98.8 % | 92.5 % | 92.4 % |
| Same residue type at matched positions | 100 % | 100 % | 100 % |
| Cα RMSD, matched (Å) | 0.075 | 0.045 | 0.085 |
| All-atom / side-chain RMSD, same residues (Å) | 0.42 / 0.61 | 0.54 / 0.75 | 0.63 / 0.89 |
| Published waters matched by ours (≤ 1 Å) | 83.9 % | 65 % | 67.4 % |
| Our waters matched by published (≤ 1 Å) | 69.5 % | 62.3 % | 62.1 % |
| Our waters in published ligand/ion sites | 15 | 7 | 5 |
| Ligands / ions in final model | GOL ×2, BME ×2, $Na^+$ ×2 | none (candidates rejected, CC < 0.70) | none (candidates rejected, CC < 0.70) |

**Exceptions and limitations**

**Supplementary Table S19. Special handling and identified problems**

| Entry / area | What happened | What was done |
|---|---|---|
| 8YYV | Deposited as one STAT1 copy plus a symmetry operator. The DNA lies on the two-fold axis in two overlapping orientations at 0.5 occupancy. A first pass refined only half the complex. | Expanded to the full dimer; each nucleotide rebuilt as alternate conformers A/B; automatic base-pair restraints turned off for this entry; baseline and refinement rerun |
| 8YR2, 9D0X, 8YS6, 9EFZ | Ligands missing from Phenix's library (PIP2, the 9D0X inhibitor A1A1I, napabucasin) or with atom names that don't match it (GABA in the published 9EFZ) | Restraints generated with eLBOW from the chemical component dictionary |
| 9C0E, 9SKT, published 9HJC | Deposited with hydrogens (46 to 7,091 atoms), one of them unknown to Phenix | Hydrogens removed; originals kept for reference |
| 8YS6 | Deposited with refined per-atom occupancies (0–1, about 560 atoms at 0) | Occupancies left unchanged, since occupancy isn't refined in this workflow |
| Coot | The rotamer fit moved backbones (broken peptides in 8Y6K and 9BW9); PDB output cut the 5-character ligand code A1LX7 to "A1L" | Side-chain-only merge added; Coot switched to mmCIF output; peptide-bond check added to every evaluation |
| Metal sites | 8YS6 $Fe_4S_4$ clusters were not coordinated to their cysteines in the deposit (Fe–Sγ > 2.8 Å) | Refinement restored Fe–Sγ at 2.26–2.27 Å; 9SKT Zn and 8Y6K Zn sites stay 4-coordinate |
| 8YR2, 11NX | Numerical failures in one polish version (Ramachandran assertion; optimizer error) | That version was dropped; the others were used |
| De novo | ModelAngelo numbered separate fragments as consecutive (9HJC A14–15, 9DTU two sites) | False-connection fix; zero-broken-peptide gate added |
